# Sex identification of ancient pinnipeds using the dog genome

**DOI:** 10.1101/838797

**Authors:** Maiken Hemme Bro-Jørgensen, Xénia Keighley, Hans Ahlgren, Camilla Hjorth Scharff-Olsen, Aqqalu Rosing-Asvid, Rune Dietz, Steven H. Ferguson, Anne Birgitte Gotfredsen, Peter Jordan, Aikaterini Glykou, Kerstin Lidén, Morten Tange Olsen

## Abstract

Determining the male and female representation in zooarchaeological material from hunted animal species is essential, to fully investigate the effects and means of prehistoric hunting practices, and may further provide valuable biological information on past animal life-history, behaviour and demography. However, the fragmented nature of the zooarchaeological record and a lack of clear diagnostic skeletal markers, often prevents such inference. Here, we test the usability of the dog nuclear genome (CanFam3.1) for sex identification of pinnipeds. First, a contemporary sample set (n=72) of ringed seals (*Pusa hispida*), with known sex was used to test the genetic sex identification method. By quantifying the proportion of X chromosome reads, as the chrX/chr1 ratio, the ratios clustered in two clearly distinguishable sex groups. Of the 72 individuals, 69 were identified to the accurate sex, which proves a high reliability of the genetic method. Second, random down sampling of a subset of the ringed seal samples to different read number, suggests at least 5000 DNA sequence reads mapped to the reference genome as the lower limit for which this method is applicable. Finally, applying this standard, sex identification was successfully carried out on a broad set of ancient pinniped samples, including walruses (*Odobenus rosmarus*), grey seals (*Halichoerus grypus*) and harp seals (*Pagophilus groenlandicus)*, which all showed clearly distinct male and female chrX/chr1 ratio groups.

## Introduction

Sex determination of ancient animal archaeological bone remains provides valuable insights to prey availability, hunting strategies and preferences of prehistoric human hunters (Weinstock, 2000a). Furthermore, it contributes with essential insight to the ecology, behaviour, demography and life history of past animal populations (Allentoft et al., 2010; Pečnerová et al., 2017). Investigating the sex ratio among prey gives an idea about their availability, hunting strategies and preferences of the human hunters (Weinstock, 2000a, b; Gotfredsen and Møbjerg 2004; Magnell 2005). Furthermore, for hunted animal species, sex determination can also be used to trace potential demographic effects of such hunting pressure on the hunted population, such as being illustrated in modern case studies (Taylor et al., 2008; Marealle et al., 2010).

Morphological sex determination of archaeological material is not always possible, as only few bone elements, and only among adult individuals, show generally accepted indicators of sex. These are specific bone characteristics of e.g. suid canines (Mayer and Brisbin, 1988), ungulate horn cores and the innominate bones (Hatting, 1995; Greenfield, 2006). It can also be sex specific size categories of certain bone measures (osteometric sex identification) (e.g. Bartosiewicz et al., 1997; Weinstock 2000b), which has been accounted for in domestic animals as well as in wild species. Furthermore, anthropogenic fragmentation prior to deposition and secondary diagenesis due to the various taphonomic processes that the skeletal remains undergo after deposition (Lyman, 1994), might alter the few morphologically sex identifiable bone parts to such a degree that sex identification is no longer reliable.

In pinnipeds, morphological sex identification is only possible for species (e.g. otariids) that, based on the size of their bones, show a pronounced sexual dimorphism between fully adult males and females. Most phocid seals, however, show only limited or no sexual dimorphism (Fay, 1985; King, 1983). Sex identification using osteometric analysis, based on measurements of the mandibles, has been established only for the walrus (Wiig et al., 2007). However, such osteometric sex identification is only possible for individuals that have reached fully adult growth. Because of the limited possibilities to apply reliable sex identification in archaeological studies, based on the morphological characteristics or osteometric analysis, it is important to seek alternative methods to investigate sex ratios.

Genetic sexing approaches have been sought as alternatives to osteological sex identification methods. For instance, some make use of the differences in the characteristic of the sex chromosomes, for which PCR primers were designed to target specific regions in which the two chromosomes differ resulting in different amplification signals (Aasen and Medrano, 1990; Faerman et al., 1995). Most methods have been developed for mammalian species e.g. focusing on the zinc-finger protein domain (Aasen and Medrano, 1990; Morin et al., 2005; Svensson et al., 2008) and the amelogenin gene (Akane et al., 1991; Faerman et al., 1995; Stone et al., 1996), both for use in, forensic studies (Akane et al., 1991), sex identification of preimplantation embryos (Aasen and Medrano, 1990), as well as for sex identification in archaeology (Faerman et al., 1995; Stone et al., 1996).

More recently, the advent of high-throughput sequencing methods allow for genetic sexing approaches beyond targeting a short specific DNA region. These methods are ideal for studies that focus on getting a broader genetic insight as sex identification can be obtained alongside of multiple other types of analyses on the same data set (Bro-Jørgensen et al., 2018). High-throughput shotgun sequencing generates a raw data set, which do not target any specific regions of the genome and therefore roughly represents the genome components of the sampled individuals. Since this is the case, it is possible to identify the sex of an individual based on the relative quantity of DNA reads representing the X chromosome (in mammals) if the DNA data yield (i.e. DNA sequence reads) is high enough. Mammalian male individuals will have a relative representation of X chromosome reads about half that of mammalian females, since males, in contrast to females, carry only one X chromosome. To carry out the sex identification, the relative quantity of DNA reads representing the X chromosome in a sample has to be determined by comparison to the autosomal chromosomes. For both males and females, the autosomal chromosomes are represented by two copies in the genome and are therefore roughly represented by reads in proportion to their chromosome size. For simplicity, as well as visual reasons, a ratio of the read representation is usually calculated based on the X chromosome and one autosomal chromosome of similar size (see for example Pečnerová et al., 2017; Bro-Jørgensen et al., 2018).

Here, we present a comparative sexing method using the annotated dog reference genome to identify the sex of a shotgun sequenced data set of ancient pinnipeds consisting of walrus (*Odobenus rosmarus*), grey seal (*Halichoerus grypus*) and harp seal (*Pagophilus groenlandicus*), based on the relative read representation of chromosome X. A data set of contemporary ringed seals (*Pusa hispida*) with known sex was used to test the accuracy of the method.

## Methods

### Sampling, DNA extraction and sequencing

#### Ancient seal and walrus samples

The bone and tooth samples obtained for ancient DNA analysis include archaeological and historical samples from walrus, grey seal and harp seal. Samples were chosen regardless of their size and level of degradation and represent in total a broad range of regions and time periods (Table S1).

The ancient seal samples were extracted using a lysis buffer consisting of EDTA (0.5M, pH8), Triton X and Proteinase K (100mg/mL). Extracts were concentrated using Amicon Ultra Centrifugal Filters (Sigma-Aldrich, Darmstadt, Germany) and eluted in MinElute-spin columns (Qiagen, Hilden, Germany). Libraries were built using the method described by Meyer and Kircher (2010). All steps were carried out in the Clean Laboratory at the Archaeological Research Laboratory, Stockholm University, Sweden. Following size selection by Ampure beads the samples were pooled and sequenced at the SciLifeLab Stockholm, Sweden. A total of 15 harp seal samples were sequenced on Illumina HiSeqX, one was sequenced on Illumina HiSeq2500 platform and the remaining 65 harp seal samples were sequenced on NovaSeq S1, while 58 of the grey seal samples were sequenced on the Illumina HiSeq2500 and one on NovaSeq S1. Further details about the grey seal samples and the laboratory work is found in Ahlgren et al. (n.d.).

The walrus samples were extracted using a lysis buffer consisting of EDTA (0.5M, pH8), Urea (1M) and Proteinase K (10mg/μL), and eluted using Zymo-spin reservoirs (Zymo Research, CA, USA) combined with MinElute-spin columns (Qiagen, Hilden, Germany) (Dabney et al., 2013). Libraries were built using the method described by Carøe et al. (2018), as detailed for walruses in Keighley et al. (2019). All steps were carried out in a dedicated ancient DNA laboratory at University of Copenhagen, Denmark.

#### Contemporary ringed seal samples

Samples from contemporary Arctic ringed seals were obtained from research monitoring programmes and Inuit subsistence hunt in Greenland and Canada. All sampled individuals were aged and sex determined in the field. DNA extractions were carried out using Thermo Scientific KingFisher Duo Prime, the KingFisher Cell and Tissue DNA Kit (Germany) and the KingFisher Duo Combi Pack for 96 DW Plate. Paired-end 150 bp libraries were built using the method described by Carøe et al. (2018), and sequencing performed using the Illumina HiSeq 4000 platform.

### Data analyses

Given a lack of published annotated pinniped genomes, we used the dog (*Canis lupus familiaris*) genome, as available on NCBI (canFam3.1), as template for assigning sex to our pinniped samples. The NCBI dog genome consists of 38 autosomal chromosomes, the X chromosome and the mitochondrial genome. As chromosome 1 is the most similar in size to chromosome X, a ratio based on the comparison of number of reads mapped to these two chromosomes were chosen for quantification of the relative quantity of DNA reads representing the X chromosome. Chromosome 1 is 122,678,785 bp long, while chromosome X is 123,869,142 bp long (NCBI resources, n.d.).

All samples were initially mapped to the dog genome using the PALEOMIX (v1.2.13.1) pipeline (Schubert et al., 2014), including SAMtools (v1.3.1) for carrying out SAM/BAM file manipulation (Li et al., 2009), AdapterRemoval (v2.2.0) (Schubert et al., 2016) and BWA (v0.7.15) for alignment and mapping of BAM-files (Li and Durbin, 2009). The ancient samples were authenticated as truly ancient by consulting the mapDamage patterns (v2.0.6) (Jónsson et al., 2013), which are characteristic for ancient DNA (Willerslev and Cooper, 2005).

Information on number of reads (hits) per chromosome for each sample were extracted from the coverage files and chrX/chr1 ratios were calculated by dividing the number of reads that mapped to chromosome X with the number of reads that mapped to chromosome 1. The total number of reads that mapped to the dog genome per sample was extracted to assess the reliability of the chrX/chr1-based sex identification.

Further, as ancient DNA analyses often are characterised by a low yield of endogenous DNA, we explored the minimum amount of DNA sequence data required to accurately determine the sex of a sample. To this end, ten male and ten female ringed seal samples were randomly sub sampled five times down to 8000, 6000, 5000, 4000, 3000 and 2000 reads, giving a total of 50 observations within each sex group for every of the six read number groups. The chrX/chr1 ratios were calculated for all observations and compared between the groups.

## Results and discussion

### Method verification on contemporary ringed seal material

The contemporary ringed seal reference samples showed an expected approximately linear increase in number of mapped reads with larger size of the chromosome (Figure 1). On this trend line, all chromosomes are represented with two copies in the genome except for chromosome X in males. Importantly, by estimating the ratio chrX/chr1 between the number of reads mapping to the X chromosome and the number of reads mapping to chromosome 1, a total of 69 out of the 72 contemporary ringed seal samples (95.8%) were assigned to the correct sex, suggesting a very high accuracy of our method. The three mismatches may owe to incorrect sex determination or reporting by the sample collectors, sample mix-up during handling and laboratory work, or less likely bias in the genetic sex determination.

**Figure 1.**
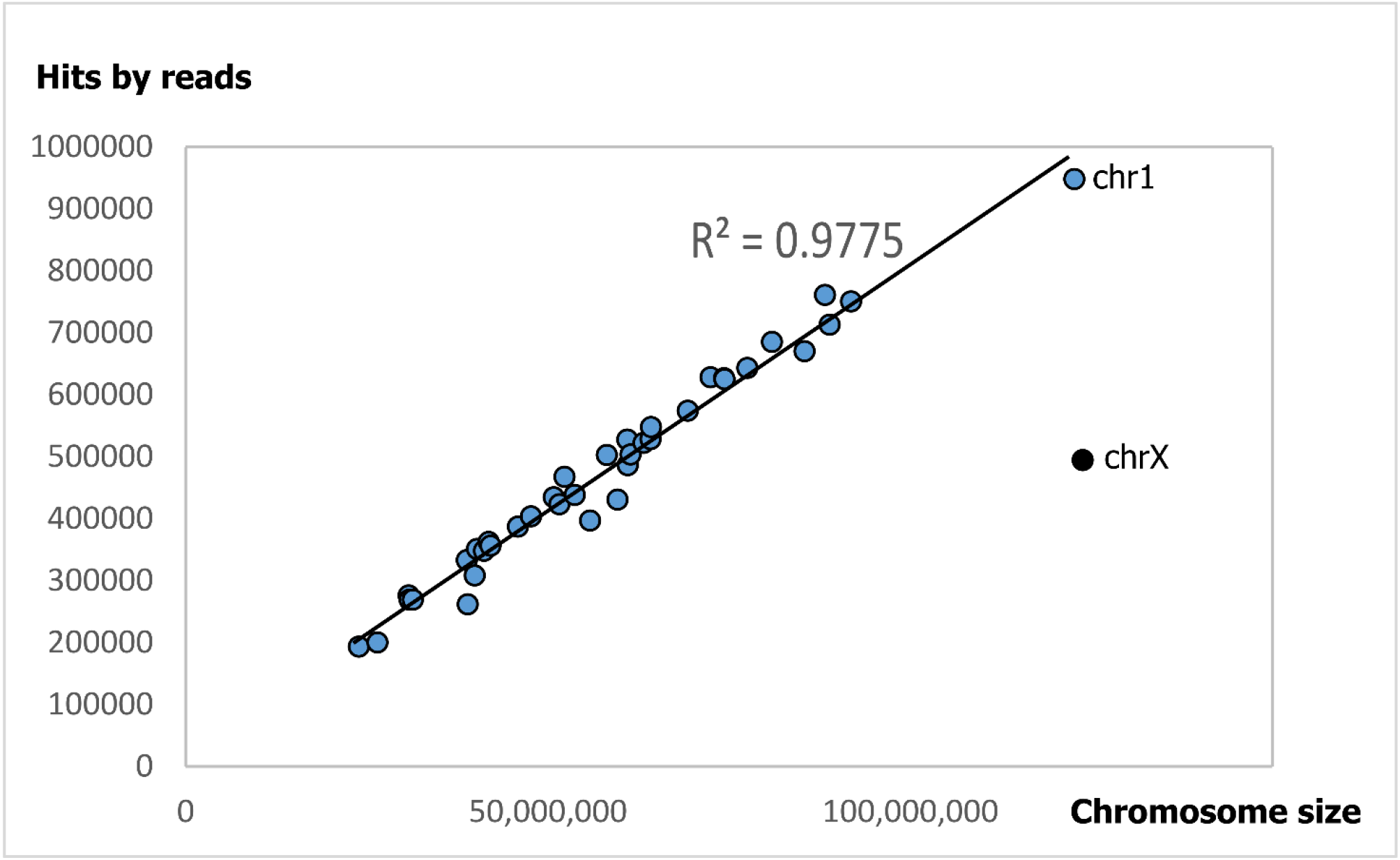
An example of a contemporary ringed seal male individual showing an approximate linear relationship between number of reads mapping to a certain autosomal chromosome (blue dots) and the size of that chromosome. Only being found in one copy in males, the chromosome X is here represented by approximately half as many mapped reads as expected by the chromosome size alone.

In the down-sampled ringed seal sequence data, there is a clear tendency that lower DNA read number leads to an increase in the standard deviation, as random variation in the down sampling will have had a bigger impact on the chrX/chr1 ratios when the read number is small (Table S2, Figure 2). The distribution is larger in the lower read groups with minor overlap between the male and female groups, while the male and female groups become more defined and distinct from each other with higher number of DNA reads. Our limited data suggest that around 5000 endogenous DNA sequence reads, at a size of roughly 150 bp, is sufficient to reliably determine the sex of a sample.

**Figure 2.**
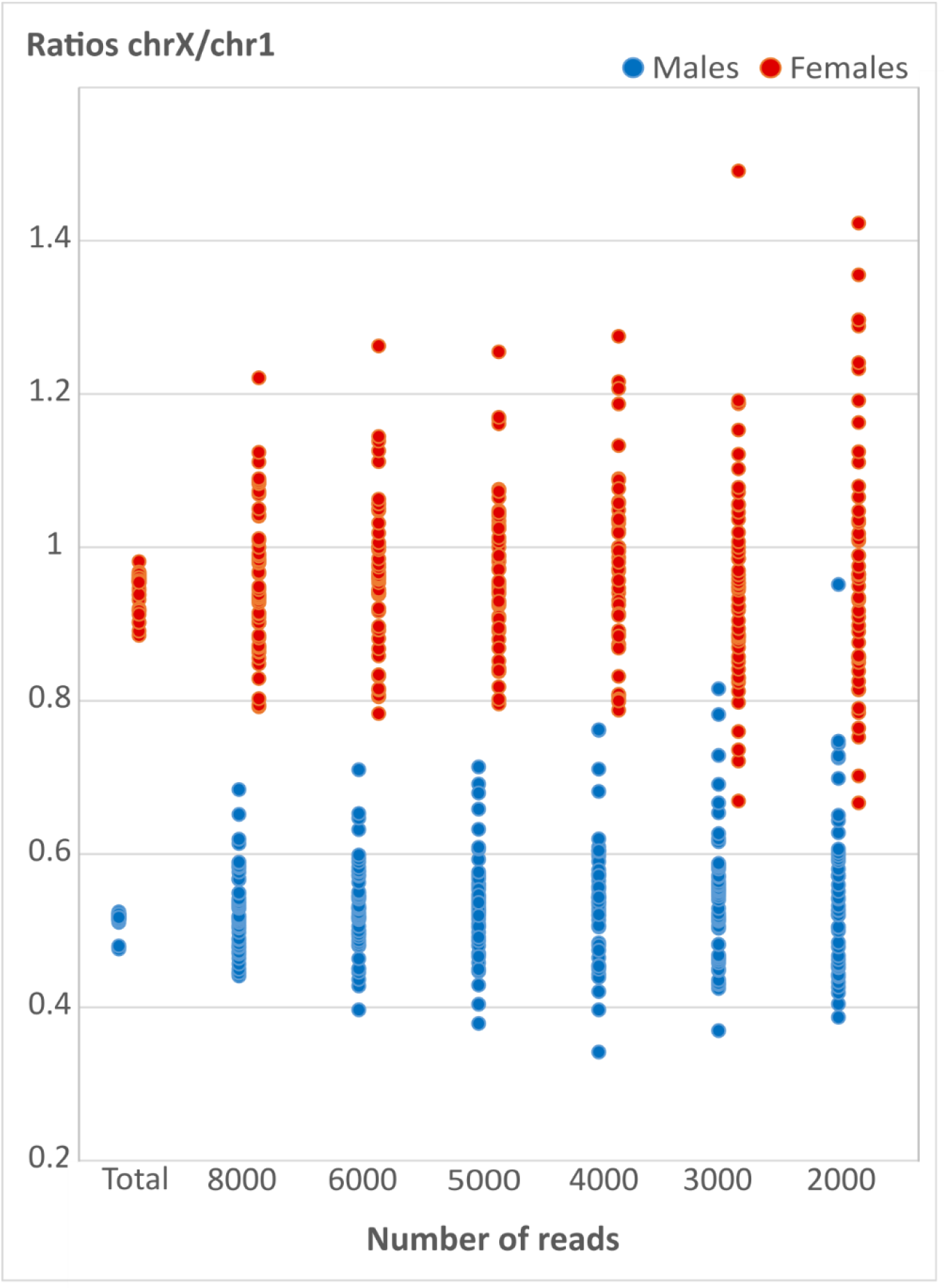
The distribution of the chrX/chr1 ratios for each of the down sampling tests as well as for the total read number of the full sample set are shown in the order of decreasing read number. Female samples are shown in red and male samples are shown in blue.

### Sex identification of ancient pinnipeds

From the initial walrus sequencing data, 41 of the 158 samples were excluded from further sex identification analysis due to too low DNA and sequence yield, with less than 100 reads mapping to the dog genome. However, for the remaining data, including the 59 grey seals and 81 harp seals, the chrX/chr1 ratio was successfully calculated for each sample, and the results evaluated based on the threshold of a minimum 5000 total number of reads mapped to the dog genome (Table 1).

**Table 1.** Ancient pinniped samples for sex identification.

| Species | Samples | Included samples (>100 reads) | Average number of reads | Sex identifiable samples (>5000 reads) | Success rate (% sex identified) | Males | Females |
| --- | --- | --- | --- | --- | --- | --- | --- |
| Walrus | 158 | 118 | 148,746.6 | 74 | 46.8% | 55 (74.3%) | 19 (25.6%) |
| Grey seal | 59 | 59 | 332,762.4 | 58 | 98.3% | 26 (44.8%) | 32 (55.2%) |
| Harp seal | 81 | 81 | 36,081.6 | 37 | 45.7% | 20 (54.1%) | 17 (45.9%) |

From a total of 298 ancient samples 169 individuals were identified to sex following these standards, giving a total sex identification success of 57% across all three pinniped species. For harp seals and walrus, a little less than 50% of the total number of samples was possible to identify to sex. The grey seal samples had an almost 100% success rate in sex identification due to extraordinary good preservation of almost all the samples (Table 1). The samples that did not satisfied the threshold of at least 5000 reads mapped to the dog reference genome, had generally low read number even before mapping.

Sorting all the chrX/chr1 ratios of the ancient samples into groups based on their total read number, illustrates the importance of using a lower threshold for sex identification. With 5000 total reads or more, no critical outliers are found in the ancient chrX/chr1 ratio distributions (Figure 3).

**Figure 3.**
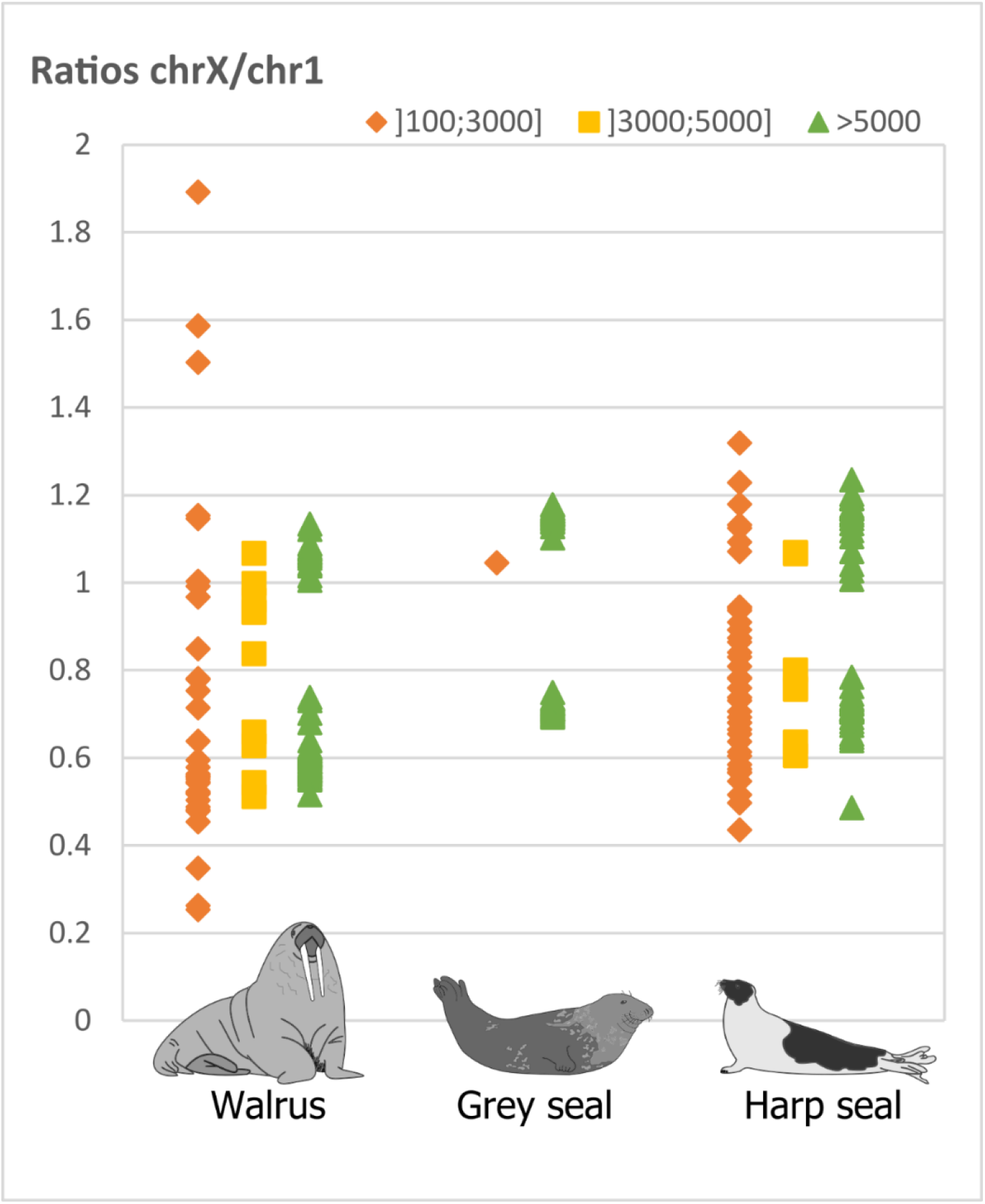
ChrX/chr1 ratios of all ancient samples which gave DNA results (>100 reads). The colours indicate the interval of the total number of reads per sample that mapped to the dog reference genome. A total of 5000 reads is presented as the limit for a reliable sex identification (Graphic design by Maiken Hemme Bro-Jørgensen).

### Is the dog genome a proper reference for pinnipeds?

In the dog, the theoretical median value of the female chrX/chr1 ratios should be close to 1.010 (123,869,142 bp divided by 122,678,785 bp), and the theoretical median value of the male chrX/chr1 ratios should be close 0.505. However, this is not the case for the pinniped samples.

Canids and pinnipeds both belong to the order of carnivore, but to different families within the caniformia suborder (Delisle and Strobeck, 2005) and their genomes differ accordingly. Most pinnipeds, including walrus, grey seal, harp seal and ringed seal, have 32 autosomal chromosomes (Arnason, 1974), while the dog genome with 38 autosomes shows quite a different karyotypic structure. Therefore, the size of chromosome 1 and X, may likely differ between dog and pinnipeds. Thus, the number of reads that will map to these two chromosomes, and hence the ratio calculated between them, might as a result differ from the theoretical value. Similarly, different chromosome sizes can be the reason for variation in the range of the chrX/chr1 values among the different pinniped species themselves (Figure 2 and 3).

The representation of the X chromosome, being about ¾ of the autosomal effective population size (Johnson and Lachance, 2012), can have had the evolutionary consequence that more affinity is found between the X chromosome of dogs and pinnipeds than is the case for the autosomes in general, thereby skewing the chrX/chr1 ratios of this study to be slightly larger than expected. Male ratios might be further skewed towards higher values due to the presence of homologous regions between the X chromosome and the Y chromosome (Lahn and Page, 1999), as a result of which Y chromosome reads are likely to have been wrongly identified as X chromosome reads.

Despite the differences in the genomes and potential issues with the representation of the X chromosome, we have shown that using the dog reference genome to quantify the relative representation of X chromosome mapping reads in pinnipeds is a valid sex identification method, as long as sufficient total read numbers are met.

### Implications for understanding human hunting practice and pinniped biology

Sex ratios in archaeological materials do not necessarily reflect the sex ratio of the ancient populations themselves. The sex ratio in archaeological assemblages depends on various ecological and anthropological parameters such as seasonal and regional prey availability, prey accessibility, the specific ecology and breeding patterns of the prey, the cultural context, hunting methods and subsistence of the hunters (Grønnow et al. 1983; Gotfredsen and Møbjerg 2004; Glykou, 2014). For hunters with a more opportunistic subsistence, the sex ratios of the hunted prey are more likely to be representative of the wild populations (Rivals et al., 2004). However, hunters might for specific reasons have hunted one sex more than the other. The different distribution of males and females e.g. in harp seals during the breeding season, might result in a higher representation of females as they are more vulnerable and thus easier to catch. For the hunter gatherers who exploited breeding colonies of harp seals in the Baltic Sea during the Mesolithic, it has been suggested that females were caught together with their pups. On the other hand, this would not be expected to be the case for regions outside the breeding areas of harp seal (Glykou, 2014). Thus, while the sex identification of this broad scale ancient zooarchaeological data set shows more or less equal proportions of males and females for grey and harp seals (Table 1), a final evaluation of these results should be embedded in the specific cultural and archaeological context. In the case of the walrus, nearly 75% of the samples were identified as male individuals (Table 1). Although our samples represent a large region and a broad time scale, it is likely that this overrepresentation of males reflect some overall degree of prey selection. Prehistoric hunting with a focus on achieving large amounts of ivory (Star et al., 2018; Pierce, 2009; Frei et al., 2015) might have primarily targeted larger dominant males, whose tusks in general are longer and thicker than the female tusks (Kastelein, 2009). While such a hunting practice would explain the skewed sex ratio, such patterns should, however, be further examined on a more fine-scaled level within each regional and cultural context. Also, taphonomic biases should be taken into account as there might be a risk that particularly male bones from species with high degree of sexual dimorphism, being larger and more robust, are more likely to have survived and being recovered during excavation. Being aware of the sampling process and including bones of varying size and level of robustness is therefore important.

Selective hunting with a preference to a specific sex can potentially cause severe problems on the population level. By targeting primarily one sex, the effect of the population loss will be the more severe as the effective population size will become even smaller. Reduction in the number of females will likely decrease the number of offspring that can be born to the next generation. Selective male hunting of polygynous species like walrus and grey seal or species with a mix between polygyny and promiscuity like harp seals (Stirlin, 1983) might not reduce the number of offspring, but will likely still over time lead to higher levels of inbreeding and risk of inbreeding depression. The risk of inbreeding depression and the effects of demographic and environmental stochasticity, particular in already small populations, can in worst case lead to the eradication of an entire population (O’Grady et al., 2006). In order to get the full picture of ancient hunting practices, the ideal is to combine sex identification with estimation of age to establish a better idea of which part of the population was being hunted.

## Conclusion

Sex identification of pinnipeds is possible using the dog reference genome for quantification of the chromosome X representation. Based on the down sampling of contemporary ringed seal samples with known sex, our results suggest that a minimum of 5000 reads in total mapped to the reference genome is required to limit the risk of samples getting incorrectly identified due to overlap between the male and the female chrX/chr1 ratio distributions. Outliers between the male and female distribution should be excluded. Due to different level of affinity to the dog reference genome, the chrX/chr1 ratios of pinnipeds might differ slightly from the theoretical values, but this will not influence the interpretation of the sex as long as the male and female ratios are clearly distinguishable. The sex identification success rates among the ancient pinniped samples were between 45.7% and 98.3% due to varying preservation states. Throughout the regions and different time periods, harp seal and grey seal showed a more or less equal sex ratio, while about 75% of the walrus samples were identified as males, which could suggest that hunting preferences has played in on the sample representation.

## Authors contributions

MHBJ, KL, and MTO conceived the study. ARA, RD and SHF provided funding, samples and sex identification of contemporary ringed seals. ABG, AG and KL identified and provide prehistoric samples. MTO and PJ provided part of the ancient samples. HA, XK, CHSO, and MHBJ carried out molecular laboratory work. MHBJ completed bioinformatics analyses. MHBJ and MTO drafted the manuscript, and all authors read, commented on and approved of the final version.

## Acknowledgements

This project has received funding from the European Union’s EU Framework Programme for Research and Innovation Horizon 2020 under Marie Curie Actions Grant Agreement No 676154 as part of “ArchSci2020”. The project was partly conducted under the BaltHealth programme, which received funding from BONUS (Art. 185), funded jointly by the EU, Innovation Fund Denmark (grants 6180-00001B and 6180-00002B), Forschungszentrum Jülich GmbH, German Federal Ministry of Education and Research (grant number FKZ 03F0767A), Academy of Finland (decision #311966) and Swedish Foundation for Strategic Environmental Research (MISTRA). We would like to thank the museums for giving us access to samples: The University Museum of Bergen, The Natural History Museum of Denmark, Greenland National Museum & Archives, The National Museum of Denmark, Archäologisches Landesmuseum in der Stiftung Schleswig-Holsteinische Landesmuseen Schloss Gottorf, Ålands Landskapsregering Museibyrån, The Swedish History Museum, The National Museum of Iceland, Icelandic Institute of Natural History, Canadian Museum of History and Canadian Museum of Nature. We would further like to thank Anne Karin Hufthammer, Ulrich Schmölcke, Kristian Gregersen, Christian Koch Madsen, Martin Appelt, Paul Szpak and Lesley Howse for procuring samples and helping with the procurement and sample authorisations. Also thanks to Fátima Sánchez Barreiro and Marta Maria Ciucani for assistance with the molecular laboratory work, and to José Alfredo Samaniego for guidance in bioinformatics.

## Supplementary material

### Supplementary tables

**Table S1.** Ancient DNA samples from pinnipeds: walrus, grey seal and harp seal. The table summarizes the total number of samples per species, the name of the collection that they belong to and the rough time periods that they represent.

| Species | Number | Collection | Time periods |
| --- | --- | --- | --- |
| Walrus | 158 | The National Museum of Denmark<br>The Natural History Museum of Denmark<br>The National Museum of Iceland<br>Icelandic Institute of Natural History<br>Canadian Museum of History<br>Canadian Museum of Nature | Pleistocene to early<br>20<sup>th</sup> century |
| Grey seal | 59 | The Swedish History Museum<br>University Museum of Bergen<br>State Museum of Schleswig | Early Mesolithic to<br>early Neolithic |
| Harp seal | 81 | The Swedish History Museum<br>University Museum of Bergen<br>State Museum of Schleswig<br>Ålands Landskapsregering Museibyrån | Late Mesolithic to<br>Bronze Age |

**Table S2.** The table presents measures of the quartiles, minimum and maximum values as well as the standard deviation of the distributions of ringed seal male and female chrX/chr1 ratios obtained from the original dataset (total) and down sampling to 8000, 6000, 5000, 4000, 3000 and 2000 reads.

| Original samples | Reads | Minimum | First quartile | Median | Third quartile | Max | SD |
| --- | --- | --- | --- | --- | --- | --- | --- |
| Ringed seal males | Total | 0.476159 | 0.515341 | 0.517535 | 0.520712 | 0.524552 | 0.011590 |
| Ringed seal females | Total | 0.885139 | 0.937784 | 0.955854 | 0.960092 | 0.981428 | 0.023279 |
| Down sampling | Reads | Minimum | First quartile | Median | Third quartile | Max | SD |
| Ringed seal males | 8000 | 0.440928 | 0.484542 | 0.518423 | 0.562465 | 0.68408 | 0.053424 |
| Ringed seal males | 6000 | 0.396721 | 0.49572 | 0.526248 | 0.560013 | 0.709779 | 0.059106 |
| Ringed seal males | 5000 | 0.378906 | 0.49967 | 0.530069 | 0.560441 | 0.713755 | 0.066876 |
| Ringed seal males | 4000 | 0.341837 | 0.489269 | 0.54258 | 0.574159 | 0.762162 | 0.081177 |
| Ringed seal males | 3000 | 0.369565 | 0.471315 | 0.54257 | 0.582605 | 0.815603 | 0.089048 |
| Ringed seal males | 2000 | 0.387097 | 0.463521 | 0.529522 | 0.593272 | 0.951807 | 0.106345 |
| Down sampling | Reads | Minimum | First quartile | Median | Third quartile | Max | SD |
| Ringed seal females | 8000 | 0.79198 | 0.889378 | 0.942057 | 1.010865 | 1.221024 | 0.093258 |
| Ringed seal females | 6000 | 0.783172 | 0.902086 | 0.959042 | 1.004673 | 1.262963 | 0.096869 |
| Ringed seal females | 5000 | 0.79562 | 0.907368 | 0.971709 | 1.028048 | 1.255319 | 0.098098 |
| Ringed seal females | 4000 | 0.787879 | 0.885193 | 0.976412 | 1.03734 | 1.275449 | 0.11569 |
| Ringed seal females | 3000 | 0.669231 | 0.871651 | 0.948116 | 1.032509 | 1.491228 | 0.141791 |
| Ringed seal females | 2000 | 0.666667 | 0.856286 | 0.96178 | 1.061355 | 1.423077 | 0.17208 |

